# Warming and altered precipitation independently and interactively suppress alpine soil microbial growth in a decadal-long experiment

**DOI:** 10.1101/2023.06.08.544195

**Authors:** Yang Ruan, Ning Ling, Shengjing Jiang, Xin Jing, Jin-Sheng He, Qirong Shen, Zhibiao Nan

**Author notes:** **Corresponding author:** Ning Ling, *E-mail address. **Corresponding author:** Jin-Sheng He, *E-mail address. **E-mail addresses of all authors** Yang Ruan, Ning Ling, Shengjing Jiang, Xin Jing, Jin-Sheng He, Qirong Shen, Zhibiao Nan.

## Abstract

Warming and precipitation anomalies affect terrestrial carbon balance partly through altering microbial eco-physiological processes (e.g., growth and death) in soil. However, little is known about how such processes responds to simultaneous regime shifts in temperature and precipitation. We used the ^18^O-water quantitative stable isotope probing approach to estimate bacterial growth in alpine meadow soils of the Tibetan Plateau after a decade of warming and altered precipitation manipulation. Our results showed that the growth of major taxa was suppressed by the single and combined effects of temperature and precipitation, eliciting 40-90% of growth reduction of whole community. The antagonistic interactions of warming and altered precipitation on population growth were common (~70% taxa), represented by the weak antagonistic interactions of warming and drought, and the neutralizing effects of warming and wet. The members in *Solirubrobacter* and *Pseudonocardia* genera had high growth rates under changed climate regimes. These results are important to understand and predict the soil microbial dynamics in alpine meadow ecosystems suffering from multiple climate change factors.

## INTRODUCTION

Global climate change is threatening multi-dimensional ecosystem services, such as terrestrial primary productivity and soil carbon storage (Jansson & Hofmockel, 2020; Walker *et al*., 2022; Zhou *et al*., 2022), especially in high-elevation ecosystems (Ma *et al*., 2017; Liu *et al*., 2018). Of these, the effects of global climate change on microbial processes related to soil carbon cycling should receive more extensive attention, because carbon balance will have feedbacks on climate system, and further reinforce/diminish the net impact on ecosystem functioning (Jansson & Hofmockel, 2020). Microbial growth and death, the critical eco-physiological processes, serve as the major engine of soil organic carbon (SOC) turnover and thus dominates the feedback on climate (Hicks *et al*., 2022; Sokol *et al*., 2022). Quantitative estimates of trait-based responses of microbes to multiple climate factors is critical for improved biogeochemical models and predicting the feedback effects to global change.

Climate warming and precipitation regime shift can influence soil microbial physiological activities directly or indirectly (Schimel, 2018; Jansson & Hofmockel, 2020; Purcell *et al*., 2022; Sokol *et al*., 2022). The Tibetan Plateau is considered among the most sensitive ecosystems to climate change (Liu *et al*., 2018). In such alpine regions, warming can alleviate low temperature limitations to enzymatic activity, stimulating SOC mineralization and microbial respiration (Dieleman *et al*., 2012; Streit *et al*., 2014). Long-term warming reduces soil organic carbon pools and exacerbates carbon limitation of soil microbes, causing a negative effect on microbial growth and eco-physiological functions (Jansson & Hofmockel, 2020; Melillo *et al*., 2017; Purcell *et al*., 2022; Streit *et al*., 2014). Precipitation fluctuation constrains microbial physiological performance and functions, which is expected to be the major consequence of future climate change in mesic grassland ecosystems (Cook Benjamin *et al*., 2015; McHugh *et al*., 2017; Oppenheimer-Shaanan *et al*., 2022; Yuan *et al*., 2017). Reduced precipitation affects soil processes notably by directly stressing soil organisms, and also altering the supply of substrates to microbes via dissolution, diffusion, and transport (Schimel, 2018). Increased frequency and magnitude of precipitation events could cause microbial species loss by ‘filtering out’ the taxa with low tolerance to increased soil moisture and drying-rewetting (Evans & Wallenstein, 2014). In addition, higher mean annual precipitation (MAP) triggers an increase in SOC decomposition (Zhou *et al*., 2022), which could cause a negative effect on microbial growth in long term. Collectively, as climate change typically causes negative consequences on the microbe-associated processes in terrestrial ecosystems.

As temperature and precipitation are of particular relevance, the interactive effects of warming and altered precipitation remain largely illusive, especially on the population growth of soil microbes (Zhu *et al*., 2016; Song *et al*., 2019). Drought limits the resistance of the entire system to warming (Hoeppner & Dukes, 2012). Higher evapotranspiration in a warmer world will result in chronically lower average soil moisture (Reich *et al*., 2018), further reducing the eco-physiological performance of soil microbes (Schimel, 2018). In contrast, enhanced precipitation relieves overall water limitations caused by warming and improved primary productivity and soil respiration (Fay *et al*., 2008). The responses of microbial population growth to multiple climate factors could be complex because i) the changed climate conditions can directly affect the eco-physiological characteristics of soil microbes and ii) indirectly affect microbial functioning by altering soil physicochemical properties (e.g., redox conditions and nutrient allocation) and aboveground plant composition (Qi *et al*., 2022; Yang *et al*., 2021). The response of decomposer growth rates to the interaction of climate factors may be strongly idiosyncratic, varying among taxa, thus making predictions at the ecosystem level difficult.

The goal of current study is to comprehensively estimate taxon-specific growth responses of soil bacteria following a decade of warming and altered precipitation manipulation on the alpine grassland of the Tibetan Plateau, by using the ^18^O-quantitative stable isotope probing (^18^O-qSIP) (Fig. 1A). We focused on the single and interactive effects of temperature (T) and precipitation (P) on the population-specific growth of soil bacteria. We classified the interaction types as additive, synergistic, weak antagonistic, strong antagonistic and neutralizing interactions between climate factors (Fig. 1B) by using the effect sizes and Hedges’ *d* (an estimate of the standardized mean difference) (Côté *et al*., 2016; Harpole *et al*., 2011; Ma *et al*., 2019; Yue *et al*., 2017). We addressed the following hypotheses: 1) long-term warming and altered precipitation regimes (i.e., drought or wet) have negative effects on microbial growth in alpine meadow soils; 2) the interactive effects between warming and altered precipitation on microbial population growth rates are not simply additive.

**Fig. 1.**
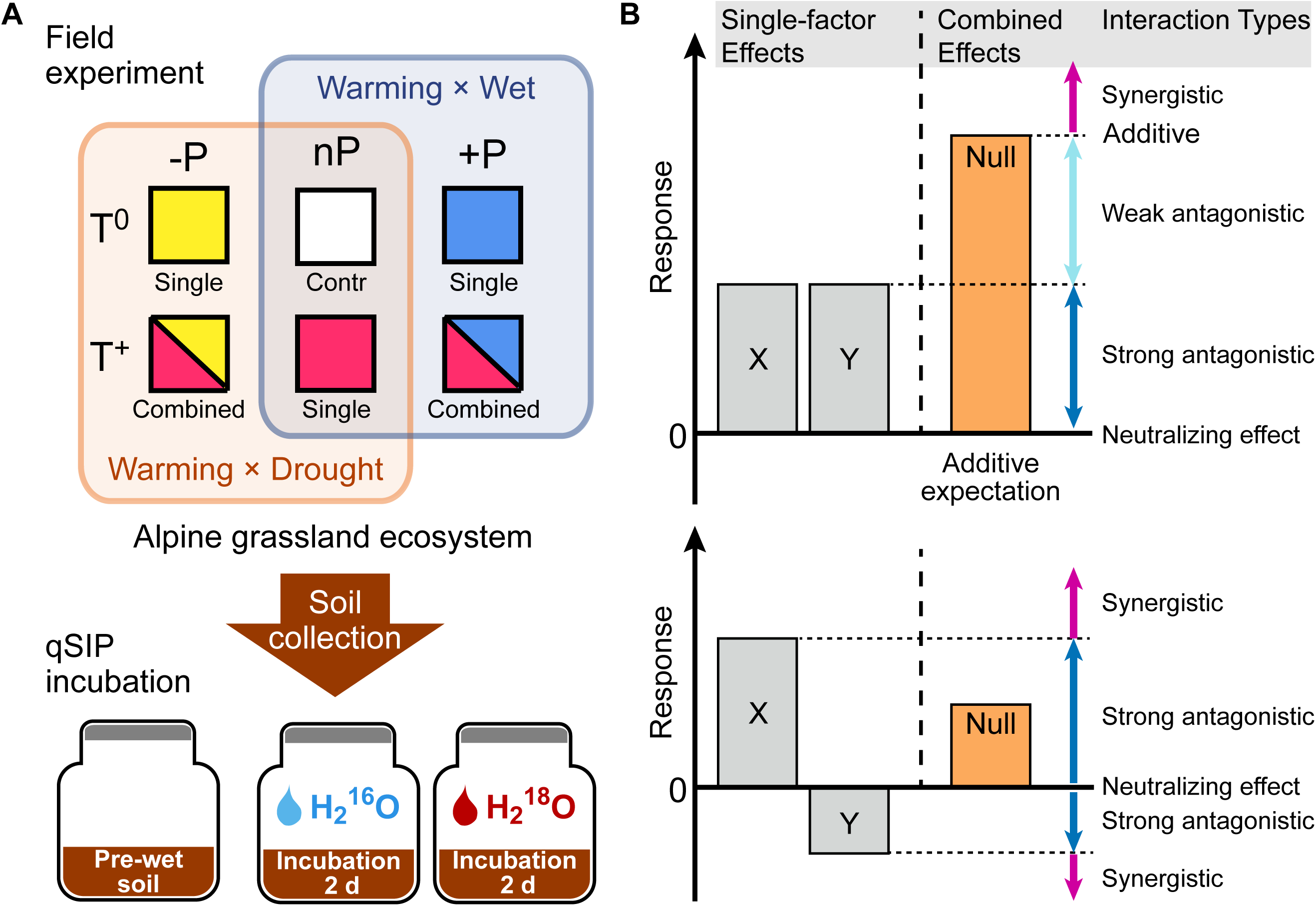
Field experiment design for simulated warming and altered precipitation, qSIP incubation, and the growth responses of soil bacteria to changing climate regimes. To examine the effects of warming and altered precipitation on an alpine grassland ecosystem, two levels of temperature (T^0^, T^+^), and three levels of precipitation (-P, nP, +P) were established in 2011. The soil samples were collected in 2020 and used for ^18^O-qSIP incubation (A). Potential interaction types between multiple climate factors (B). The diagram shows that two factors (X and Y) of warming and altered precipitation impact a biological response in the same direction (upper) or have opposing effects on when acting separately. Their combined effect could be additive, i.e. the sum of the two factor effects. Alternatively, the interaction types can be antagonistic or synergistic. Null model (we use the additive expectation as the null model here) provides the threshold for distinguishing between these interactions.

## MATERIALS AND METHODS

### Study design and soil sampling

The warming-by-precipitation experiment was established in 2011 at the Haibei National Field Research Station of Alpine Grassland Ecosystem (37°37′N, 101°33′E, with elevation 3215 m), which is located on the northeastern Tibetan Plateau in Qinghai Province, China. The climate type is a continental monsoon with mean annual precipitation of 485 mm and the annual average temperature approximately −1.7°C. The high rainfall and temperature mainly occur in the peak-growing season (from July to August (Liu *et al*., 2018)). The soils are Mat-Gryic Cambisols, with the average pH value of surface soil (0-10 cm) being 6.4 (Ma *et al*., 2017).

The experimental design has been described previously in (Ma *et al*., 2017). Briefly, experimental plots were established in an area of 50 m × 110 m in 2011, using a randomized block design with warming and altered precipitation treatments. Each block contained six plots (each plot was 1.8 m × 2.2 m), crossing two levels of temperature [ambient temperature (T^0^), elevated temperature of top 5 cm layer of the soil by 2°C (T^+^)], and three levels of precipitation [natural precipitation (nP, represents ambient condition), 50% reduced precipitation (-P, represents ‘drought’ condition) and 50% enhanced precipitation (+P, represents ‘wet’ condition)]. In the warming treatments, two infrared heaters (1,000 mm length, 22 mm width) were suspended in parallel at 150 cm above the ground within each plot. Rain shelters were used to control the received precipitation in the experimental plots. Four ‘V’-shaped transparent polycarbonate resin channels (Teijin Chemical, Japan) were fixed at a 15° angle, above the infrared heaters, to intercept 50% of incoming precipitation throughout the year. The collected rainfall from the drought plots was supplied to the wet plots manually after each precipitation event by sprinklers, increasing precipitation by 50%. To control for the effects of shading caused by infrared heaters, two ‘dummy’ infrared heaters and four ‘dummy’ transparent polycarbonate resin channels were installed in the control plots. Each treatment had six replicates, resulting in thirty-six plots.

Soil samples for qSIP incubation were collected in August 2020. Considering the cost of qSIP experiment (i.e., the use of isotopes and the sequencing of a large number of DNA samples), we randomly selected three out of the six plots, serving as three replicates for each treatment. In each plot, three soil cores of the topsoil (0-5 cm in depth) were randomly collected and combined as a composite sample, which can be considered as a mixture of rhizosphere and bulk soils. Each sampling point was as far away from infrared heaters as possible to minimize the impact of physical shading on the plants. The fresh soil samples were shipped to the laboratory and sieved (2-mm) to remove root fragments and stones.

### ^18^O-qSIP incubation

We estimated the population-specific growth rates of active microbes by conducting a ^18^O-water incubation experiment combined with DNA quantitative stable isotope probing (Fig. 1A). The incubations were similar to those reported in a previous study (Ruan *et al*., 2023). Soil samples of ambient temperature treatments (including T^0^-P, T^0^nP, and T^0^+P) were air-dried at 14 °C (average temperature across the growth season), while the soil samples of warming treatments (including T^+^-P, T^+^nP, and T^+^+P) were air-dried at 16 °C (increased temperature of 2 °C). There is no violent air convection in the incubator and the incubation temperature is relatively low (compared to 25°C used in previous studies), resulting slower evaporation and no significant discoloration caused by severe soil dehydration after 48 h. A portion of the air-dried soil samples was taken as the pre-wet treatment (i.e., before incubation without H_2_O addition). We incubated the air-dried soils (2.00 g) with 400 μl of 98 atom% H_2_^18^O (^18^O treatment) or natural abundance water (^16^O treatment) in the dark for 2 d by using sterile glass aerobic culture bottles (Diameter: 29 mm; Height: 54 mm). After incubation, soils were destructively sampled and stored at −80°C immediately. A total of 54 soil samples, including 18 pre-wet samples (6 treatments × 3 replicates) and 36 incubation samples (6 treatments × 3 replicates × 2 types of H_2_O addition), were collected.

### DNA extraction and isopycnic centrifugation

Total DNA from all the collected soil samples was extracted using the FastDNA™ SPIN Kit for Soil (MP Biomedicals, Cleveland, OH, USA) according to the manufacturer’s instructions. Briefly, the mechanical cell destruction was attained by multi-size beads beating at 6 m s^-1^ for 40 s, and then FastDNA™ SPIN Kit for Soil (MP Biomedicals, Cleveland, OH, USA) was used for DNA extraction. All DNA samples were extracted by the same person within 2-3 hours, and a unifying procedure of cell lysis and DNA extraction was used. The concentration of extracted DNA was determined fluorometrically using Qubit® DNA HS (High Sensitivity) Assay Kits (Thermo Scientific™, Waltham, MA, USA) on a Qubit® 4 fluorometer (Thermo Scientific™, Waltham, MA, USA). The DNA samples of 2-d incubation were used for isopycnic centrifugation, according to a previous publication (Ruan *et al*., 2023). Briefly, 3 μg DNA were added into 1.85 g ml^-1^ CsCl gradient buffer (0.1 M Tris-HCl, 0.1 M KCl, 1 mM EDTA, pH = 8.0) with a final buoyant density of 1.718 g ml^-1^. Approximately 5.1 ml of the solution was transferred to an ultracentrifuge tube (Beckman Coulter QuickSeal, 13 mm × 51 mm) and heat-sealed. All tubes were spun in an Optima XPN-100 ultracentrifuge (Beckman Coulter) using a VTi 65.2 rotor at 177,000 g at 18 °C for 72 h with minimum acceleration and braking.

Immediately after centrifugation, the contents of each ultracentrifuge tube were separated into 20 fractions (~250 μl each fraction) by displacing the gradient medium with sterile water at the top of the tube using a syringe pump (Longer Pump, LSP01- 2A, China). The buoyant density of each fraction was measured using a digital hand-held refractometer (Reichert, Inc., Buffalo, NY, USA) from 10 μl volumes. Fractionated DNA was precipitated from CsCl by adding 500 μl 30% polyethylene glycol (PEG) 6000 and 1.6 M NaCl solution, incubated at 37 °C for 1 h and then washed twice with 70% ethanol. The DNA of each fraction was then dissolved in 30 μl of Tris-EDTA buffer.

### Quantitative PCR and Sequencing

Total 16S rRNA gene copies for DNA samples of all the fractions were quantified using the primers for V4-V5 regions: 515F (5′-GTG CCA GCM GCC GCG G-3′) and 907R (5′-CCG TCA ATT CMT TTR AGT TT-3′) (Guo *et al*., 2018). The V4-V5 primer pairs were chosen to facilitate integration and comparison with data from previous studies (Ruan *et al*., 2023; Zhang *et al*., 2016). Plasmid standards were prepared by inserting a copy of purified PCR product from soil DNA into *Escherichia coli*. The *E. coli* was then cultured, followed by plasmid extraction and purification. The concentration of plasmid was measured using Qubit DNA HS Assay Kits. Standard curves were generated using 10-fold serial dilutions of the plasmid. Each reaction was performed in a 25-μl volume containing 12.5 μl SYBR Premix Ex Taq (TaKaRa Biotechnology, Otsu, Shiga, Japan), 0.5 μl of forward and reverse primers (10 μM), 0.5 μl of ROX Reference Dye II (50 ×), 1 μl of template DNA and 10 μl of sterile water. A two-step thermocycling procedure was performed, which consisted of 30 s at 95°C, followed by 40 cycles of 5 s at 95°C, 34 s at 60°C (at which time the fluorescence signal was collected). Following qPCR cycling, melting curves were conducted from 55 to 95°C with an increase of 0.5°C every 5 s to ensure that results were representative of the target gene. Average PCR efficiency was 97% and the average slope was −3.38, with all standard curves having R^2^ ≥ 0.99.

The DNA of pre-wet soil samples (unfractionated) and the fractionated DNA of the fractions with buoyant density between 1.703 and 1.727 g ml^-1^ (7 fractions) were selected for 16S rRNA gene sequencing by using the same primers of qPCR (515F/907R). The fractions with density between 1.703 and 1.727 g ml^-1^ were selected because they contained more than 99% gene copy numbers of each sample. A total of 270 DNA samples [18 total DNA samples of prewet soil + 252 fractionated DNA samples (6 treatments × 3 replicates × 2 types of water addition × 7 fractions)] were sequenced using the NovaSeq6000 platform (Genesky Biotechnologies, Shanghai, China).

The raw sequences were quality-filtered using the USEARCH v.11.0 (Edgar, 2010). In brief, the paired-end sequences were merged and quality filtered with “fastq_mergepairs” and “fastq_filter” commands, respectively. Sequences < 370 bp and total expected errors > 0.5 were removed. Next, “fastx_uniques” command was implemented to identify the unique sequences. Subsequently, high-quality sequences were clustered into operational taxonomic units (OTUs) with “cluster_otus” commandat a 97% identity threshold, and the most abundant sequence from each OTU was selected as a representative sequence. The taxonomic affiliation of the representative sequence was determined using the RDP classifier (version 16) (Wang *et al*., 2007). In total, 19,184,889 reads of the bacterial 16S rRNA gene and 6,938 OTUs were obtained. The sequences were uploaded to the National Genomics Data Center (NGDC) Genome Sequence Archive (GSA) with accession numbers CRA007230.

### Quantitative stable isotope probing calculations

As ^18^O labeling occurs during cell growth via DNA replication, the amount of ^18^O incorporated into DNA was used to estimate the growth rates of active taxa. The density shifts of OTUs between ^16^O and ^18^O treatments were calculated following the qSIP procedures (Hungate *et al*., 2015; Koch *et al*., 2018). Briefly, the number of 16S rRNA gene copies per taxon (e.g., genus or OTU) in each density fraction was calculated by multiplying the relative abundance (acquisition by sequencing) by the total number of 16S rRNA gene copies (acquisition by qPCR). Then, the GC content and molecular weight of a particular taxon were calculated. Further, the change in ^18^O isotopic composition of 16S rRNA genes for each taxon was estimated. We assumed an exponential growth model over the course of the incubations, and average population growth rates within 2-d incubation were estimated. The absolute rate of growth is a function of the rate of appearance of ^18^O-labeled 16S rRNA genes. Therefore, the growth rate of taxon *i* was calculated as:

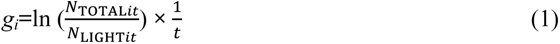

where N_TOTAL*it*_ is the number of total gene copies for taxon *i* and N_LIGHT*it*_ represents the unlabeled 16S rRNA gene abundances of taxon *i* at the end of the incubation period (time *t*). N_LIGHT*it*_ is calculated by a function with four variables: N_TOTAL*it*_, average molecular weights of DNA (taxon *i*) in the ^16^O treatment (M_LIGHT*i*_) and in the ^18^O treatment (M_LAB*i*_), and the maximum molecular weight of DNA that could result from assimilation of H_2_^18^O (M_HEAVY*i*_) (Koch *et al*., 2018). We further calculated the average growth rates (represented by the production of new16S rRNA gene copies of each taxon per g dry soil per day) along the incubation, using the following equation (Stone *et al*., 2021):

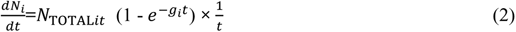

where *t* is the incubation time (day). Please see the supplementary material for detailed qSIP calculation principles and procedures. All data calculations were performed using the qSIP package (https://github.com/bramstone/qsip) in R (Version 3.6.2) (Team 2006).

### Single and combined effects of climate change factors

To address the effects of warming and altered precipitation on microbial growth rates, three single-factor effects (warming, 50% reduced precipitation only, and 50% enhanced precipitation only) and two combined effects (combined warming and reduced precipitation manipulation and combined warming and enhanced precipitation manipulation) were calculated by the natural logarithm of response ratio (lnRR), representing the response of microbial growth rates in the climate change treatment compared with that in the ambient treatment (Yue *et al*., 2017). The lnRR for growth rates was calculated as:

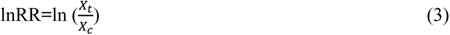

where *X_t_* is the observed growth rates in climate treatment and *X_c_* is that in control. Three lnRR values (i.e., three replicates) were calculated, and 95% confidence interval (CI) was estimated using a bootstrapping procedure with 1000 iterations (Ruan *et al*., 2023). If the 95% CI did not overlap with zero, the effect of treatment on microbial growth is significant.

### The interaction between warming and altered precipitation

Briefly, all six climate treatments were divided into two groups, warming combined with reduced precipitation scenario (Warming × Drought), and warming combined with enhanced precipitation scenario (Warming × Wet), by using the ambient temperature and precipitation treatment (T^0^nP) as control (Fig. 1A). Hedges’ *d*, an estimate of the standardized mean difference, was used to assess the interaction effects of warming × drought (i.e. reduced precipitation) and warming × wet (i.e. enhanced precipitation), respectively (Yue *et al*., 2017). The interaction effect size (*d*_I_) of warming × drought or warming × wet was calculated as:

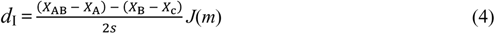

where *X*_c_, *X*_A_, *X*_B_ and *X*_AB_ are one of three duplicate values of growth rates in the control, treatment groups of factor A, B, and their combination (AB), respectively. Total three *d*_I_ values were calculated in each comparison, and 95% CI was estimated using a bootstrapping procedure with 1,000 iterations. The *s* and *J(m)* are the pooled standard deviation and correction term for small sample bias, respectively, which were calculated by the following equations:

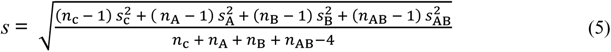

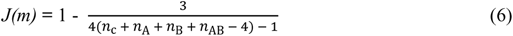

where *n*_c_, *n*_A_, *n*_B_ and *n*_AB_ are the sample sizes, and *s*_c_, *s*_A_, *s*_B_ and *s*_AB_ are the standard deviations in the control, experimental groups of A, B, and their combination (AB), respectively.

The interaction types between warming and altered precipitation were mainly classified into three types, i.e. Additive, Synergistic and Antagonistic, according to the single-factor effects and 95% CI of *d*_I_. If the 95% CI of *d*_I_ overlapped with zero, the interactive effect of warming and altered precipitation was additive. The synergistic interaction included two cases: 1) the upper limit of 95% CI of *d*_I_ < 0 and the single-factor effects were either both negative or have opposite directions; 2) the lower limit of 95% CI of *d*_I_ > 0 and both single-factor effects were positive. The antagonistic interaction also included two cases: 1) the upper limit of 95% CI of *d*_I_ < 0 and both single-factor effects were positive; 2) the lower limit of 95% CI of *d*_I_ > 0 and the single-factor effects were either both negative or have opposite directions (Yue *et al*., 2017). We further divided antagonistic interaction into three sub-categories: weak antagonistic interaction, strong antagonistic interaction, and neutralizing effect, by comparing the single-factor and combined effect sizes (Fig. 1B). The weak antagonistic interaction determined if the combined effect size was larger than the single-factor effect sizes, but smaller than their expected additive effect. The strong antagonistic interaction determined if the combined effect size was smaller than the single-factor effect sizes but not equal to zero. The neutralizing effect represented the combined effect size is equal to zero, and at least one single-factor effect size is not equal to zero.

### Statistical analyses

Uncertainty of growth rates (95% CI) was estimated using a bootstrapping procedure with 1,000 iterations (Ruan *et al*., 2023). The cumulative growth rates at the phylum-level were estimated as the sum of taxon-specific growth rates of those OTUs affiliated to the same phylum. Significant differences of bacterial growth rates for each group between climate treatments were assessed by two-way ANOVA in R (version 3.6.2). Phylogenetic trees were constructed in Galaxy /DengLab (http://mem.rcees.ac.cn:8080) with PyNAST Alignment and FastTree functions (Caporaso *et al*., 2009; Price *et al*., 2009). The trees were visualized and edited using iTOL (Letunic & Bork, 2016). To estimate the phylogenetic patterns of incorporators whose growth subjected to different factor interaction types, the nearest taxon index (NTI) was calculated by the “picante” package in R (version 3.6.2) (Webb *et al*., 2002). NTI with values larger than 0 and their *P* values less than 0.05 represent phylogenetic clustering. The *P* values of NTI between 0.05 and 0.95 represent random phylogenetic distributions. KO gene annotation of taxa was performed by PICRUSt2 (Phylogenetic Investigation of Communities by Reconstruction of Unobserved States), which predicted functional abundances based on marker gene sequences (Douglas *et al*., 2020). The marker genes related to carbon (C), nitrogen (N), sulfur (S), and phosphorus (P) cycling were selected according to the conclusions reported in previous documents (Dai *et al*., 2019; Llorens-Marès *et al*., 2015; Nelson Albright *et al*., 2015).

## RESULTS

### Overall growth response of soil bacteria to warming and altered precipitation

Excess atom fraction ^18^O value (Fig. 2) and the population growth rate of each OTU were calculated using the qSIP pipeline. Collectively, 1373 OTUs were identified as “^18^O incorporators” (i.e. OTUs with growth rates significantly greater than zero) and used for subsequent data analyses. The maximum cumulative growth rates of the whole communities occurred in the ambient temperature and ambient precipitation condition (T^0^nP), and all climate manipulations had negative effects on soil bacterial growth (Fig. 3A). The individual impact of warming, drought, and wet conditions resulted in the most substantial negative effects on bacterial growth compared with the combined effects of warming × drought and warming × wet. A result that illustrates the antagonistic interactions between warming and modified precipitations patterns (Fig. 3B). Moreover, the combined effect size of wet and warming was smaller than that of drought and warming, indicating a higher degree of antagonism of warming × wet.

**Fig. 2.**
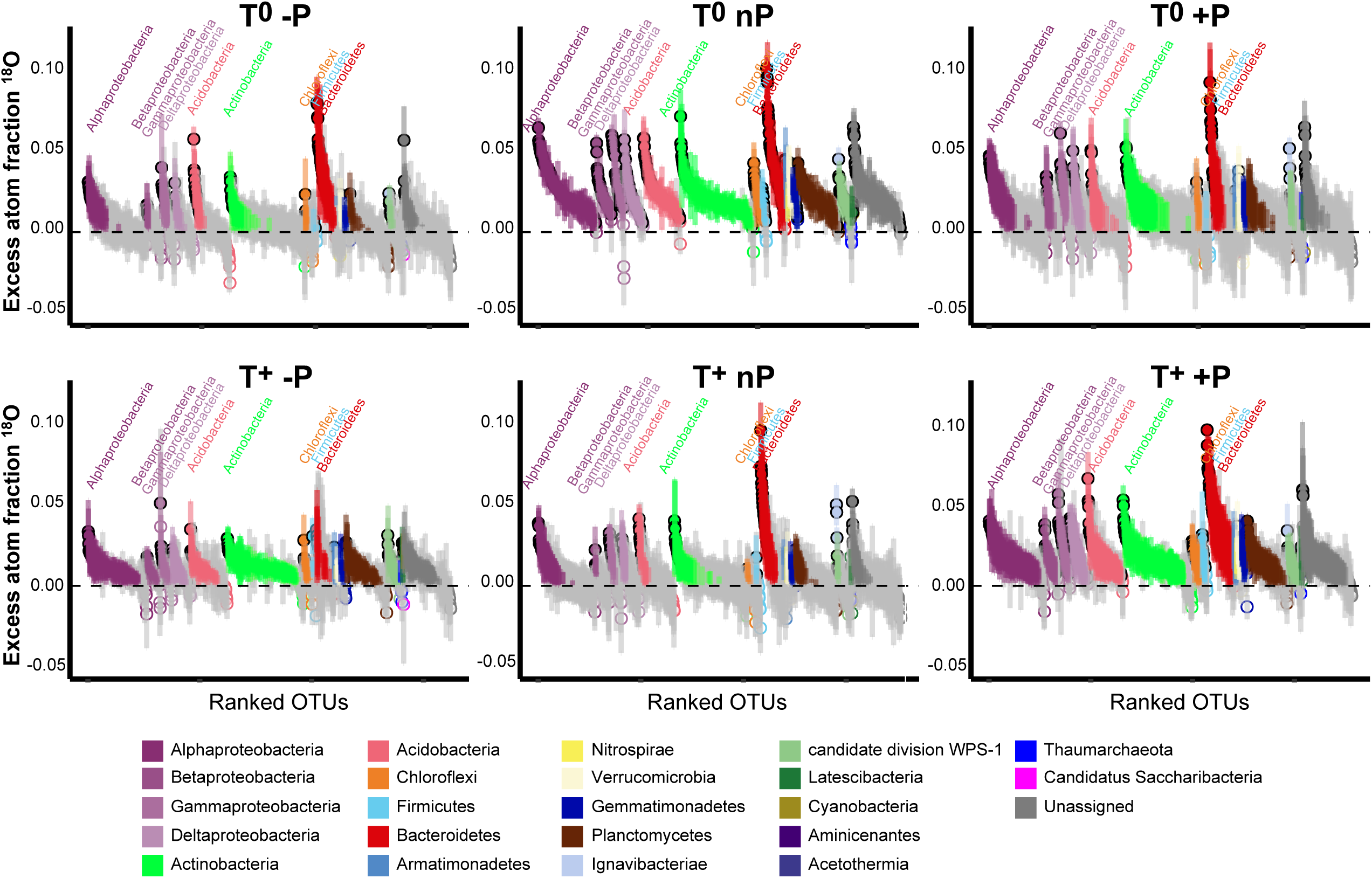
Species-specific shifts of ^18^O excess atom fraction (EAF-^18^O). Bars represent 95% confidence intervals (CIs) of OTUs. Each circle represents an OTU and color indicates phylum. The open circles with gray bars represent OTUs with 95% CI intersected with zero (indicating no significant ^18^O enrichment); Closed circles represent the OTUs enriched ^18^O significantly, whose 95% CIs were away from zero (i.e., the OTUs had detectable growth).

**Fig. 3.**
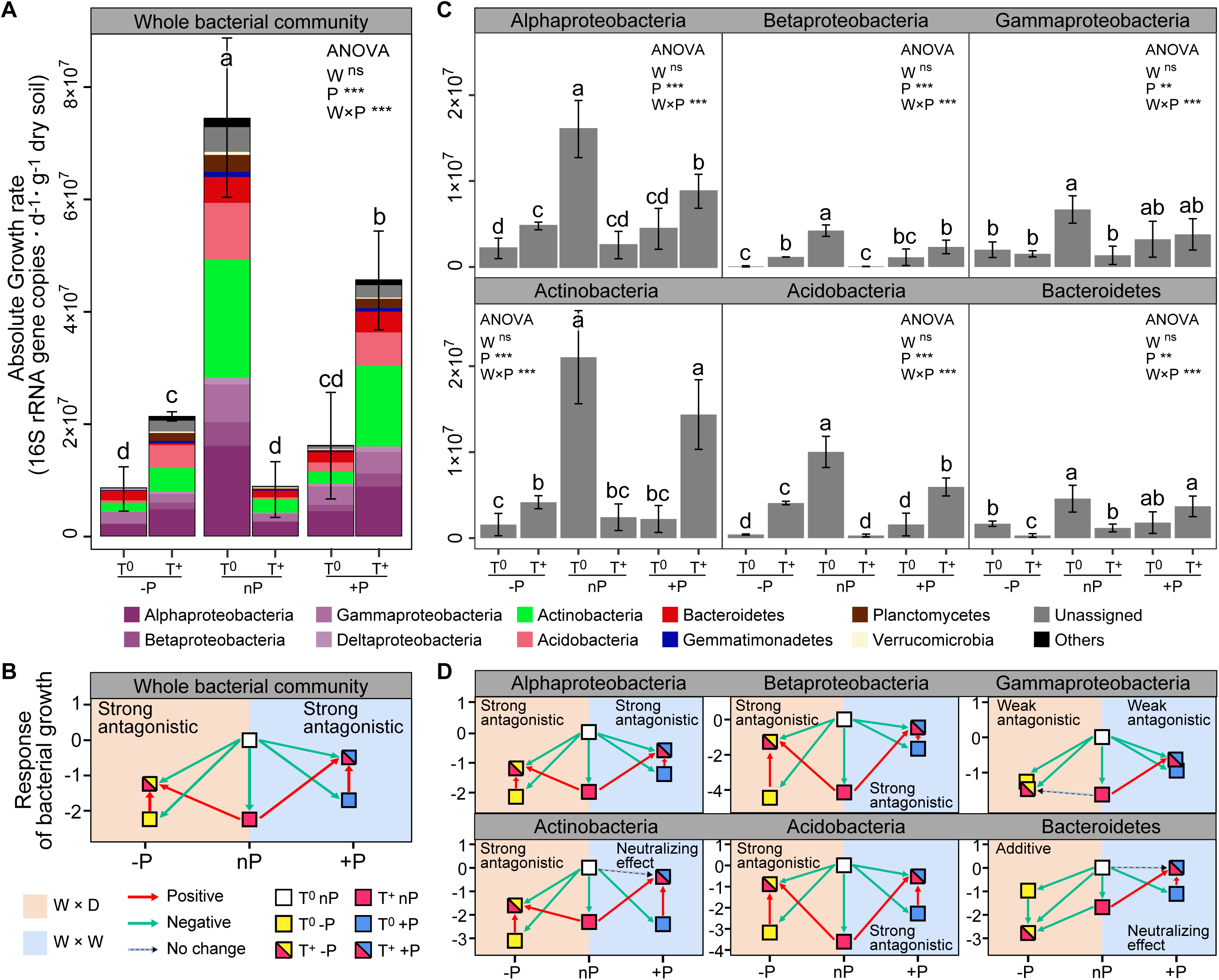
Bacterial growth responses to climate change and the interaction types between warming and altered precipitation. The growth rates (**A**), and responses of soil bacteria to warming and altered precipitation (**B**) at the whole community level. The growth rates (**C**), and responses of the dominant bacterial phyla (**D**) had similar trends with that of the whole community. Values represent mean and the error bars represent standard deviation. Different letters indicate significant differences between climate treatments.

Growth of the major bacterial phyla was also negatively influenced by the individual climate factors (Fig. 3C and 3D). The antagonistic interactions of T × P were prevalent among the major phyla (except Bacteroidetes showed the additive interaction between drought and warming). We also found the significant smaller combined effect sizes of warming × wet in the major phyla compared with that of warming × drought (*P* < 0.05), such as Actinobacteria, Bacteroidetes and Betaproteobacteria, indicating higher degree of antagonism. In Actinobacteria and Bacteroidetes, the effect of wet and warming neutralized each other, as the combined effect of these two factors had no effect on growth.

### Phylogeny for the species whose growth subjected to different factor interactions

We constructed a phylogenetic tree including all ^18^O incorporators in all six climate treatments (Fig. 4A). The single-factor effects on the growth of incorporators tended to be negative (Fig. 4B): Warming (T^+^nP) reduced the growth of 75% of the taxonomic groups, which was followed by drought and wet (74% and 67%, respectively). Warming × drought and warming × wet had the smaller impacts on the growth of incorporators, compared with the single effects (especially T^+^+P, only 43% of incorporators showed negative growth responses). The interaction type of T × P on the growth of ~70% incorporators was antagonistic (i.e., the combined effect size is smaller than the additive expectation) (Fig. 4C). The weak antagonistic interaction on bacterial growth was dominant under the warming × drought conditions (41% of incorporators), while more incorporators (34%) whose growth subjected to neutralizing effect was found under the warming × wet conditions. These findings were robust at a subOTUs level by the zero-radius OTU (ZOTU) analysis (Fig. S1 and S2).

**Fig. 4.**
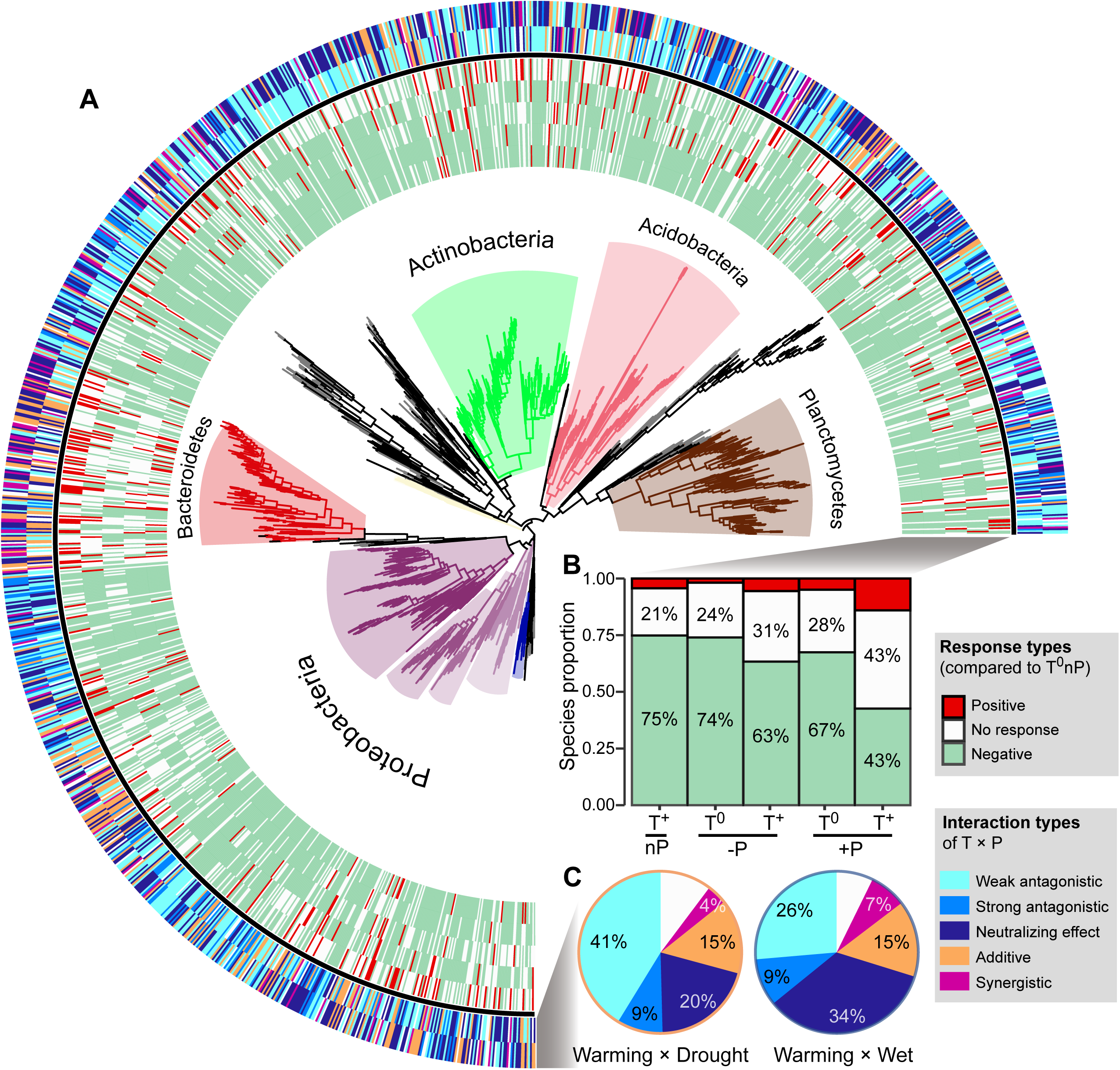
The growth responses and phylogenetic relationship of incorporators subjected to different interaction types under two climate scenarios. A phylogenetic tree of all incorporators observed in the grassland soils (**A**). The inner heatmap represents the single and combined factor effects of climate factors on species growth, by comparing with the growth rates in T^0^nP. The outer heatmap represents the interaction types between warming and altered precipitation under two climate change scenarios. The proportions of positive or negative responses in species growth to single and combined manipulation of climate factors by summarizing the data from the inner heatmap (**B**). The proportions of species growth influenced by different interaction types under two climate change scenarios by summarizing the data from the outer heatmap (**C**).

Phylogenetic relatedness can provide information on the ecological and evolutionary processes that influenced the emergence of the eco-physiological responses in taxonomic groups (Evans & Wallenstein, 2014). Nearest taxon index (NTI) was used to determine whether the species in a particular growth response are more phylogenetically related to one another than to other species (i.e., close or clustering on phylogenetic tree) (Table S1). NTI is an indicator of the extent of terminal clustering, or clustering near the tips of the tree (Evans & Wallenstein, 2014; Webb *et al*., 2002). Overall, the most incorporators whose growth was influenced by the antagonistic interaction of T × P showed significant phylogenetic clustering (i.e., species clustered at the phylogenetic branches; NTI > 0, *P* < 0.05). The incorporators whose growth subjected to the additive interaction of warming × drought also showed significant phylogenetic clustering (*P* < 0.05), but randomly distributed under warming × wet scenario (*P* = 0.116). In addition, incorporators whose growth is influenced by the synergistic interaction of T × P showed random phylogenetical distribution under both climate scenarios (*P* > 0.05).

### Higher degree of antagonism in warming and wet scenario

We further assigned the antagonistic intensity to the five interaction types on a 5-point scale, from −1 to 3 for synergistic, additive, weak antagonistic, strong antagonistic and neutralizing effect, respectively (Fig. S3A), where the larger values represent higher degree of antagonism. Then, the overall antagonistic intensities of all incorporators under warming × drought and warming × wet scenarios were estimated by weighting the relative proportions of incorporators subjected to different interaction types (Fig. S3B). We found higher overall antagonistic intensity of warming × wet than that of warming × drought, contributing by a higher proportion of incorporators whose growth subjected to neutralizing effect (Fig. 4C and Fig. S3B).

Of the total 1373 incorporators, 1218 were shared in both warming × drought and warming × wet scenarios (Fig. 5A). That is, the difference in interactive effects between warming × drought and warming × wet we observed was due to a within-species change in growth response, rather than changes in species composition. Of these species identified in both warming × drought and warming × wet scenarios, 453 were assigned a higher degree of antagonistic interaction of warming × wet than that of warming × drought. Further, the growth of 215 incorporators were influenced by the weak antagonistic interaction of warming × drought, and neutralizing effect of warming × wet. The growth response of these 215 species could contribute mainly to the overall growth patterns observed in grassland bacterial community under warming and altered precipitation scenarios, because of the prevalence of weak antagonistic interaction of warming × drought and neutralizing effect of warming × wet (Fig. 4C).

**Fig. 5.**
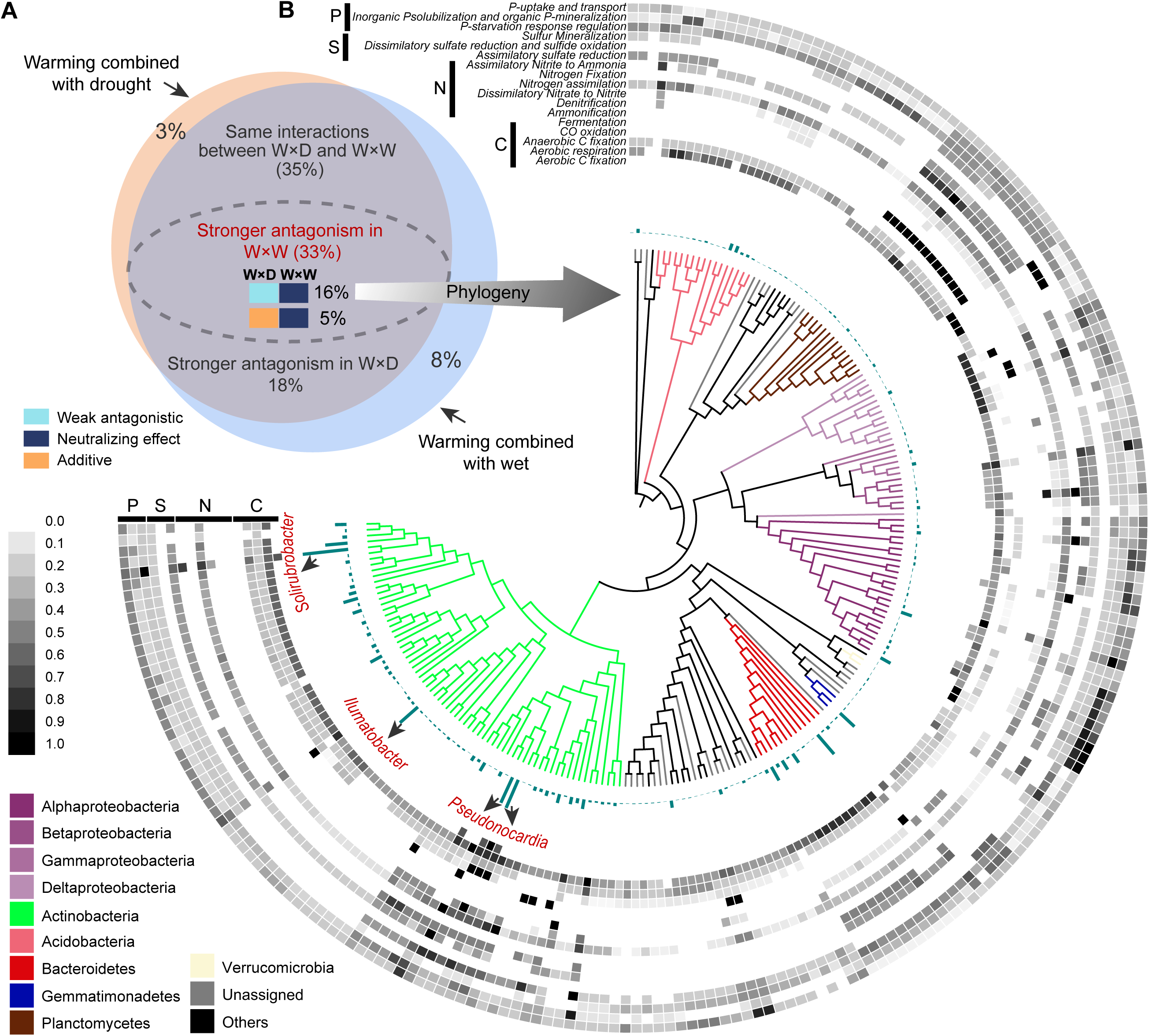
Within-species shift in interaction types contributed to the variance of the whole community growth response under two climate scenarios. Venn plots showing the overlaps of incorporators, and their interaction types between two climate scenarios (**A**). The phylogenetic relationship of the 215 incorporators whose growth dynamics were influenced by the weak antagonistic interaction of warming × drought and by the neutralizing effect of warming × wet (**B**). The blue-green bars represent the average growth rates of incorporators across different climate treatments. The heatmap displayed the potential functions associated with carbon and nutrient cycles predicted by Picrust2. The values of function potential were standardized (range: 0-1). The “anta” represents the antagonistic interaction; “W×D” represents warming × drought and “W×W” represents warming × wet.

We further assessed the potential functional traits of these 215 incorporators subjected to the dominant interaction types by PICRUST2 software (Fig. 5B). The top three OTUs with the highest growth rates possessed extremely high species abundance (Supplementary Data 1). The three taxa also possessed a higher functional potential: the member affiliated to *Solirubrobacter* (OTU 14), has the high functional potential for aerobic C fixation and CO oxidation, nitrogen assimilation and assimilatory nitrite to ammonia, and phosphatase synthesis and phosphate transport transport-related functions. The members affiliated to the genus *Pseudonocardia* (OTU 5 and OTU 31), harbor a higher functional potential for aerobic C fixation, aerobic respiration, and CO oxidation, dissimilatory nitrate to nitrite and nitrogen assimilation, and sulfur mineralization functions. Furthermore, we annotated the genomic characteristics by aligning species sequences to the GTDB database (Genome Taxonomy Database), and we found that OTU 14 (*Solirubrobacter*) was predicted to have larger genomes and proteomes (Supplementary Data 1). All these results suggested that these three species could play essential roles at the species and functional levels of ecosystems.

## DISCUSSION

Microbial populations might respond differently to environmental changes, resulting in differential contributions to ensuing biogeochemical fluxes (Blazewicz *et al*., 2020). Here, we estimated microbial growth responses by using the qSIP technique to decadal-long warming and altered precipitation regimes in the alpine grassland ecosystem on the Tibetan Plateau, which is considered highly susceptible and vulnerable to climate change (Ma *et al*., 2017). After a decade of temperature and precipitation regime shift, the pervasive negative impacts of a single climate factors on soil bacterial growth in alpine grassland ecosystem were observed (Fig. 3), which supports our first hypothesis that long-term warming and altered rainfall events consistently reduce microbial growth. Consistent with our findings, a recent experimental study demonstrated that 15 years of warming reduced the growth rate of soil bacteria in a montane meadow in northern Arizona (Purcell *et al*., 2022). These negative effects of climate factors on microbial growth are likely due to the variation related to availability of soil moisture and organic carbon (Dieleman *et al*., 2012; Wu *et al*., 2011). Previous evidences suggest that warming has a negative impact on soil carbon pools (Jansson & Hofmockel, 2020; Purcell *et al*., 2022), mainly because of the rapid soil carbon mineralization and respiration (Melillo *et al*., 2017). Carbon is the critical element in cell synthesis, the reduction of microbially accessible carbon pools may explain the diminished microbial growth after long-term warming. In addition, long-term warming can induce soil water deficiency (Dieleman *et al*., 2012; Jansson & Hofmockel, 2020), thereby slowing microbial growth.

Altered rainfall patterns that result in increased aridity or wetter conditions, mediate ecosystem cycling by affecting above- and below-ground biological processes (Song *et al*., 2019). As soil water availability is reduced, changes in osmotic pressure cause microbial death or dormancy, while others can accumulate solutes to survive under lower water potentials (Schimel, 2018). However, such accumulation of osmolytes could depend on highly energetic expenses (Boot *et al*., 2013; Jansson & Hofmockel, 2020; Schimel *et al*., 2007), resulting in less energetic allocation to growth (trade-offs between microbial growth and physiological maintenance). On the other hand, intensified rainfall patterns alter the composition and life strategies of soil bacteria, enriching the taxa with higher tolerance to frequent drying-rewetting cycles (Evans & Wallenstein, 2014). Such taxa may possess physiological acclimatization, such as synthesizing chaperones to stabilize proteins and thicker cell wall to withstand osmotic pressure (Schimel *et al*., 2007). These adaptation and acclimation strategies also create physiological costs (Schimel *et al*., 2007), increasing carbon allocation to physiological maintenance instead of new biomass (Lipson, 2015).

Climate-induced changes in the growth and structure of plant communities can also influence soil microbial growth by altering the amount and quality of plant-derived carbon (Bardgett *et al*., 2013). Increasing drought reduced the transfer of recently fixed plant carbon to soil bacteria and shifts the bacterial community towards slow growth and drought adaptation (Fuchslueger *et al*., 2014). A 17-year study of California grassland provided evidence that terrestrial net primary production (NPP) to precipitation gradient are hump-shaped, peaking when precipitation is near the multi-year mean growing season level (Zhu *et al*., 2016). Reduced NPP could affect plant carbon inputs to the soil, ultimately having a negative effect on microbial growth.

Characterizing the interactive effects of multiple global change drivers on microbial physiological traits is important for predicting ecosystem responses and soil biogeochemical processes (Song *et al*., 2019; Zhu *et al*., 2016). In this study, a decade-long experiment revealed that bacterial growth in alpine meadows is primarily influenced by the antagonistic interaction between T × P (Fig. 3 and 4). A range of ecosystem processes have been revealed to be potentially subject to antagonistic interactions between climate factors, for instance, net primary productivity (Shaw *et al*., 2002), soil C storage and nutrient cycling processes (Dieleman *et al*., 2012; Wu *et al*., 2011; Larsen *et al*., 2011). Reduced precipitation can mute organic carbon mineralization by inhibiting soil respiration, which could maintain a relatively adequate soil carbon content and explain the diminished negative effects on microbial growth by the combined manipulation of warming and drought (Jansson & Hofmockel, 2020; Wu *et al*., 2011). Conversely, enhanced precipitation could stimulate SOM decomposition, causing further loss of soil carbon under warming conditions (Zhou *et al*., 2022). However, increased rainfall can also alleviate the moisture limitation on plant growth induced by warming, increasing plant-derived carbon inputs (Jansson & Hofmockel, 2020; Wu *et al*., 2011). The increased carbon inputs may alleviate microbial carbon limitation in soil, which partly explains the higher microbial growth rates under the combined treatment of warming and enhanced precipitation relative to single climate factor manipulation.

The degree of phylogenetic relatedness can indicate the processes that influenced community assembly, like the extent a community is shaped by environmental filtering (clustered by phylogeny) or competitive interactions (life strategy is phylogenetically random distribution) (Evans & Wallenstein, 2014; Webb *et al*., 2002).The results showed that the incorporators whose growth was influenced by the antagonistic interaction of T × P showed significant phylogenetic relatedness, indicating the occurrence of taxa more likely shaped by environment filtering (i.e., selection pressure caused by changes in temperature and moisture conditions). In contrast, the growing taxa affected by synergistic interactions of T × P showed random phylogenetic distributions (Table S1), which may be explained by competition between taxa with similar eco-physiological traits or changes in genotypes (possibly through horizontal gene transfer) (Evans & Wallenstein, 2014). We also found that the extent of phylogenetic relatedness to which taxa groups of T × P interaction types varied by climate scenarios, suggesting that different climate history processes influenced the ways bacteria survive temperature and moisture stress.

About one-third of bacterial species had growth with higher levels of antagonistic interaction of warming × wet than that of warming × drought (Fig. 5A). By annotating the genomic information, we further found that the species with the high growth rate (OTU 14, *Solirubrobacter*) has a relatively larger genome size and protein coding density (Supplementary Data 1), indicating larger gene and function repertoires. A previous study showed that the genus *Solirubrobacter* detected in the Thar desert of India is involved in multiple biochemical processes, such as N and S cycling (Kunjukrishnan Kamalakshi *et al*., 2018). Members in the genus *Solirubrobacter* are also considered to contribute positively to plant growth (Liu *et al*., 2020), and can be used to predict the degradation level of grasslands, indicating the critical roles on maintaining ecosystem services (Yan *et al*., 2022). This is, however, still to be verified, as the functional output from PICRUSt2 is less likely to resolve rare environment-specific functions (Douglas *et al*., 2020). This suggests the development of methods combining qSIP with metagenomes and metatranscriptomes to assess the functional shifts of active microorganisms under global change scenarios. Note that the experimental parameters such as DNA extraction and PCR amplification efficiencies also have significant effects on the accuracy of growth assessment. This alerts the need to standardize experimental practices to ensure more realistic and reliable results.

The evaluation of ecosystem models based on results obtained from single-factor experiments usually overestimate or underestimate the impact of global change on ecosystems, because the interactions between climate factors may not be simply additive (Dieleman *et al*., 2012; Wu *et al*., 2011; Zhou *et al*., 2022). Our results demonstrated that both warming and altered precipitation negatively affect the growth of grassland bacteria; However, the combined effects of warming and altered precipitation on the growth of ~70% soil bacterial taxa were smaller than the single-factor effects, suggesting antagonistic interaction. This suggests the development of multifactor manipulation experiments in precise prediction of future ecosystem services and feedbacks under climate change scenarios.

## ACKNOWLEDGEMENTS

This work was supported by the National Science Foundation of China [42277100 (NL)].

## Statement of authorship

NL contributed the conceptualization and supervision of the experiment; YR conducted the experiment and data analysis, and wrote original draft; SJJ and JSH provided samples; QRS, ZBN and JSH contributed to project scope; And all authors contributed substantially to the review and editing of manuscript.

## Data accessibility statement

The sequence data were uploaded to the National Genomics Data Center (NGDC) Genome Sequence Archive (GSA) with accession numbers CRA007230.

