## Supplementary Materials for "Warming and altered precipitation independently and interactively suppress alpine soil microbial growth in a decadal-long experiment"

Yang Ruan *et al.*

**This PDF file includes:**

Supplementary Methods

Fig. S1-S3

Table S1

### Supplementary Methods

#### Data processing of 16S rRNA gene sequencing for zero-radius OTU (ZOTU) analysis

The sequences were quality-filtered using the USEARCH v.11.0. The paired-end sequences were merged and then quality filtered with “fastq_mergepairs” and “fastq_filter” commands, respectively. Sequences < 370 bp and total expected errors > 0.5 were removed. Next, “fastx_uniques” command was implemented to identify the unique sequences. Denoising attempts to identify all correct biological sequences in the reads, which is done by the “unoise3” command. A denoised sequence is called a "ZOTU" (zero-radius OTU).

#### Calculating taxon-specific changes in density

Taxon-specific changes in density caused by isotope incorporation were calculated as shown in equations 1 to 12 below. We calculated the total number of 16S rRNA gene copies ($y_{ijk}$) for bacterial taxon $i$ in density fraction $k$ of replicate $j$ as follows:

$$y_{ijk} = p_{ijk} \times f_{jk} \tag{1}$$

where $p_{ijk}$ is the relative abundance of each individual taxon ($i$) within an individual density fraction ($k$) from a particular replicate tube ($j$). $f_{jk}$ is the total number of 16S rRNA gene copies using the universal 16S rRNA primer for qPCR of each fraction ($k$) in each replicate density gradient ($j$).

The total number of 16S rRNA gene copies ($y_{ij}$) for bacterial taxon ($i$) in replicate ($j$) is summed across all $K$ density fractions as follows:

$$y_{ij} = \sum_{1}^{K} y_{ijk} \tag{2}$$

The density ($W_{ij}$) for bacterial taxon ($i$) of replicate ($j$) was computed as a weighted average, summing across all $K$ density fractions the density ($x_{jk}$) of each individual fraction times the total number of 16S rRNA gene copies ($y_{ijk}$) in that fraction, expressed as a proportion of the total 16S rRNA gene copies ($y_{ij}$) for taxon ($i$) in replicate ($j$), as follows:

$$W_{ij} = \sum_{1}^{K} x_{jk} \times \left(\frac{y_{ijk}}{y_{ij}}\right) \quad (3)$$

For a given taxon ($i$), we calculated the difference in density caused by isotope incorporation ($Z_i$) as follows:

$$Z_i = W_{LABi} \times W_{LIGHTi} \quad (4)$$

where $W_{LABi}$ and $W_{LIGHTi}$ are the mean, across all replicates, of the $^{18}$O-labeled treatment and unlabeled treatment.

### Calculating taxon-specific GC content and molecular weight

The GC content of each bacterial taxon ($G_i$) was calculated using the mean density for the unlabeled ($W_{LIGHTi}$) treatments. Below is the linear relationship between GC content ($G_i$, expressed as a proportion) and unlabeled buoyant density ($W_{LIGHTi}$) on a CsCl gradient (derived from pure cultures of microbial species with known and strongly differing GC contents) according to Hungate et al. (2015):

$$G_i = \frac{1}{0.083506} \times (W_{LIGHTi} - 1.646057) \quad (5)$$

The natural abundance molecular weight of DNA is a function of GC content, based on the atomic composition of the four DNA nucleotides. Single-stranded DNA made of pure adenine (A) and thymine (T) has an average molecular weight of 307.691 g/mol. The corresponding average molecular weight for DNA comprising only guanine (G) and cytosine (C) is 308.187 g/mol. When theGCcontent is known, the average molecular weight of a single strand of DNA can be calculated using the following equation:

$$M_{LIGHTi} = 0.496 G_i + 307.691 \quad (6)$$

There are 12 oxygen atoms per DNA nucleotide pair, regardless of GC content: 6 each for G and C, 7 for T, and 5 for A. These atoms contain $^{18}O$ at natural abundance, which we assume to be 0.002000429 atom fraction for $^{18}O$ (Hungate et al. 2015). The maximum labeling is achieved when all oxygen atoms are replaced by $^{18}O$. Therefore, given the molecular weight of each additional neutron (1.008665 g/mol; Hungate et al. 2015), the maximal increase in molecular weight (corresponding to 1 atom fraction $^{18}O$, or 100% atom percent $^{18}O$) is 12.07747 g/mol. The theoretical maximum molecular weight ($M_{\mathrm{HEAVYMAX}i}$) of fully $^{18}O$ -labeled DNA for taxon ($i$) is then calculated as follows:

$$M_{\mathrm{HEAVYMAX}i} = 12.07747 + M_{\mathrm{LIGHT}i} \quad (7)$$

However, as oxygen in DNA is derived from both water and organic sources, we also estimated the maximum molecular weight of DNA that could result from assimilation of $^{18}O$-water ($M_{\mathrm{HEAVY}i}$). This enables a more accurate estimate of growth, by accounting for the proportion of oxygen atoms in newly synthesized DNA derived from the labeled environmental water ($U$):

$$M_{\mathrm{HEAVY}i} = 12.07747U + M_{\mathrm{LIGHT}i} \quad (8)$$

$U$ was estimated via sensitivity analysis, with logical lower and upper bounds. Values of $U$ were rejected as too low if they resulted in a value for fully labeled DNA that was less than the molecular weight of labeled DNA observed in the 18O treatment ($M_{\mathrm{HEAVY}i} < M_{\mathrm{LAB}i}$), and values of U were rejected as too high if they resulted in estimates of unlabeled 16S rRNA gene copies at the end of the incubation that exceeded the measured abundance at the beginning of the incubation ($N_{\mathrm{LIGHT}it} > N_{\mathrm{LIGHT}i0}$), a violation of the assumption that all newly formed DNA in the 18O treatment was isotopically labeled. For all taxa, we used the empirical value of $U$ (0.60) according to (Koch et al., 2018).

The molecular weight of DNA for taxon ($i$) in the labeled treatment ($M_{\mathrm{LAB}i}$) was calculated as follows:

$$M_{\mathrm{LAB}i} = \left(\frac{z_i}{W_{LIGHTi}} + 1\right) \times M_{LIGHTi} \quad (9)$$

The atom fraction excess of $^{18}O$ for taxon ($i$) ($A_{\text{OXYGEN}i}$), accounting for the background fractional abundance of $^{18}O$ (0.002000429; Hungate et al. 2015), is then calculated as follows:

$$A_{\text{OXYGEN}i} = \left(\frac{M_{\text{LAB}i} - M_{\text{LIGHT}i}}{M_{\text{HEAVYMAX}i} - M_{\text{LIGHT}i}}\right) \times (1\text{-}0.002000429) \qquad (10)$$

### Modeling taxon-specific population growth

To estimate taxon-specific bacterial growth, we used the model of $^{18}O$ isotope substitution in DNA including an exponential model of population growth (Koch et al., 2018; Stone et al., 2021). We used a mixing model of DNA molecular weight to estimate abundances of unlabeled and labeled DNA fragments containing copies of 16S rRNA genes.

For each bacterial taxon ($i$), we assumed that the abundance of cells at time $t$ was proportional to the abundance of 16S rRNA gene copies ($N_{\text{TOTAL}i}$, with units of 16S rRNA gene copies/g soil). We further assumed that changes in bacterial abundances followed an exponential growth model over the 2-d incubation period, with the net rate of population growth ($r_i$, units: $d^{-1}$):

$$N_{\text{TOTAL}it} = N_{\text{TOTAL}i0} \times e^{r_i t} \qquad (11)$$

At time 0, the total abundance of 16S rRNA gene copies ($N_{\text{TOTAL}i0}$) was equivalent to the abundance of unlabeled 16S rRNA gene copies at the beginning of the incubation ($N_{\text{TOTAL}i0}$), but by the end of the incubation period, both unlabeled ($N_{\text{LIGHT}it}$) and labeled ($N_{\text{HEAVY}it}$) 16S rRNA gene copies may have been present such that:

$$N_{\text{TOTAL}it} = N_{\text{LIGHT}it} + N_{\text{HEAVY}it} \qquad (12)$$

Taxon-specific abundances of 16S rRNA gene copies at the beginning ($N_{\text{TOTAL}i0} = N_{\text{LIGHT}i0}$) and the end of the incubation ($N_{\text{TOTAL}it}$) were calculated as the products of the total abundance of 16S rRNA gene copies across all taxa, determined by qPCR, and the relative abundance of 16S rRNA gene copies associated with taxon i, determined by sequencing. We used a linear mixing model of DNA molecular weights to estimate the abundance of labeled 16S rRNA gene copies at the end of the incubation ($N_{\text{HEAVY}it}$), and by difference, the abundance of unlabeled 16S rRNA gene copies at the end of the incubation ($N_{\text{LIGHT}it}$):

$$N_{\mathrm{LIGHT}it} = N_{\mathrm{TOTAL}it} \times \left(\frac{M_{\mathrm{HEAVY}i} - M_{\mathrm{LAB}i}}{M_{\mathrm{HEAVY}i} - M_{\mathrm{LIGHT}i}}\right) \tag{13}$$

where for each taxon ($i$), $N_{\mathrm{TOTAL}it}$ is the total abundance of 16S rRNA gene copies at the end of the incubation, $M_{\mathrm{HEAVY}i}$ is the molecular weight of $^{18}O$-labeled DNA, $M_{\mathrm{LIGHT}i}$ is the molecular weight of unlabeled DNA in $^{16}O$ treatment, and $M_{\mathrm{LAB}i}$ is the average molecular weight of DNA at the end of the $^{18}O$-$H_2O$ incubation (i.e., $^{18}O$ treatment).

Then, the growth rate of taxon ($i$) was calculated as:

$$g_i = \ln \left(\frac{N_{\mathrm{TOTAL}it}}{N_{\mathrm{LIGHT}it}}\right) \times \frac{1}{t} \tag{14}$$

where $N_{\mathrm{TOTAL}it}$ is the number of total gene copies for taxon i and $N_{\mathrm{LIGHT}it}$ represents the unlabeled 16S rRNA gene abundances of taxon ($i$) at the end of the incubation period (time $t$).

We further calculated the average growth rates (represented by the production of new16S rRNA gene copies of each taxon per g dry soil per day) along the incubation, using the following equation (Stone et al., 2021):

$$\frac{dN_i}{dt} = N_{\mathrm{TOTAL}it} \left(1 - e^{-g_i t}\right) \times \frac{1}{t} \tag{15}$$

where $t$ is the incubation time (day).

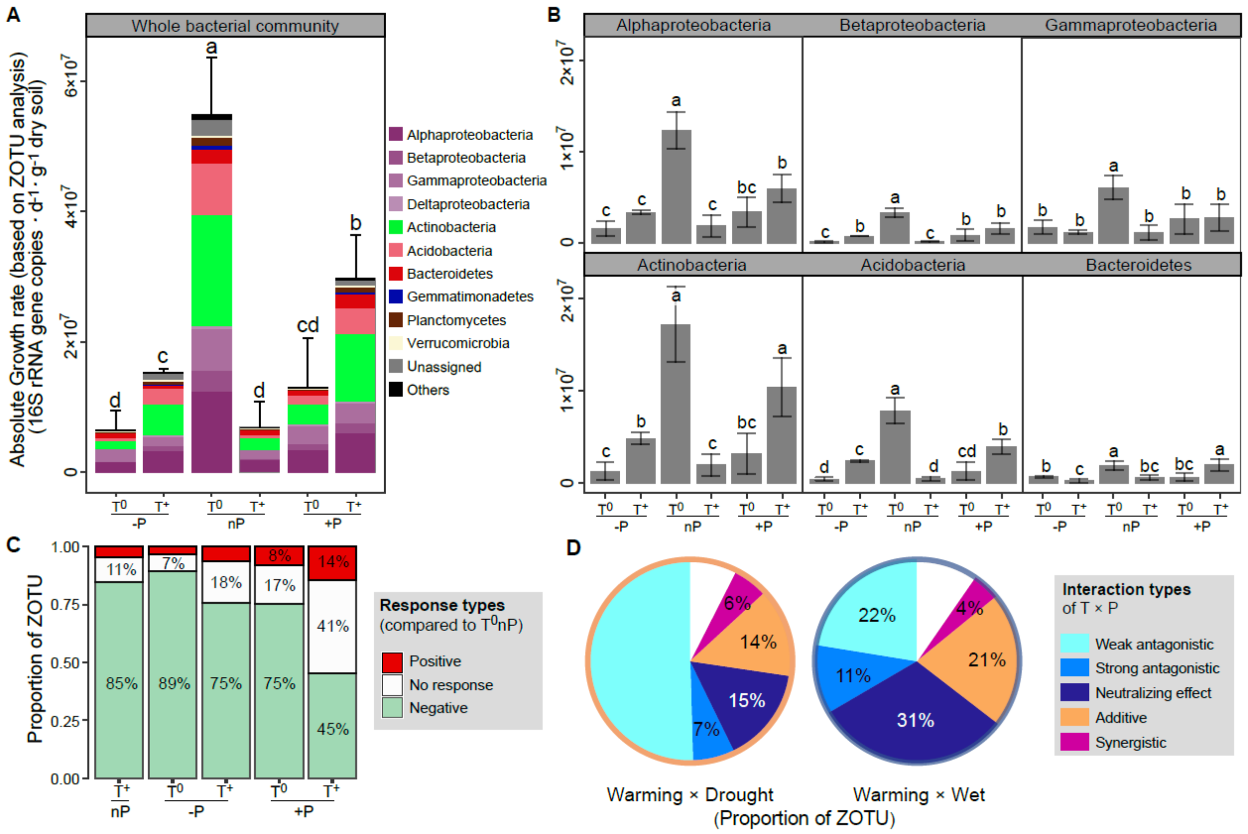


**Fig. S1.**

**The growth responses of grassland bacteria to warming and altered precipitation based on ZOTU analysis.** The results of growth rates at the community level (**A**), the phylum level (**B**), and the ZOTU level (**C** and **D**) were similar to those based on OTU analysis. **C** the single and combined factor effects of climate factors on species growth, by comparing with the growth rates in $T^0$nP. **D** the proportions of species growth influenced by different interaction types of T × P. $T^0$-P represents the ambient temperature and decreased precipitation; $T^+$-P represents warming and decreased precipitation; $T^0$cP represents ambient temperature and precipitation; $T^+$cP represents warming and ambient precipitation; $T^0$+P represents ambient temperature and enhanced precipitation; $T^+$+P represents warming and enhanced precipitation. Values represent mean and the error bars represent standard deviation. Different letters indicate significant differences between climate treatments.

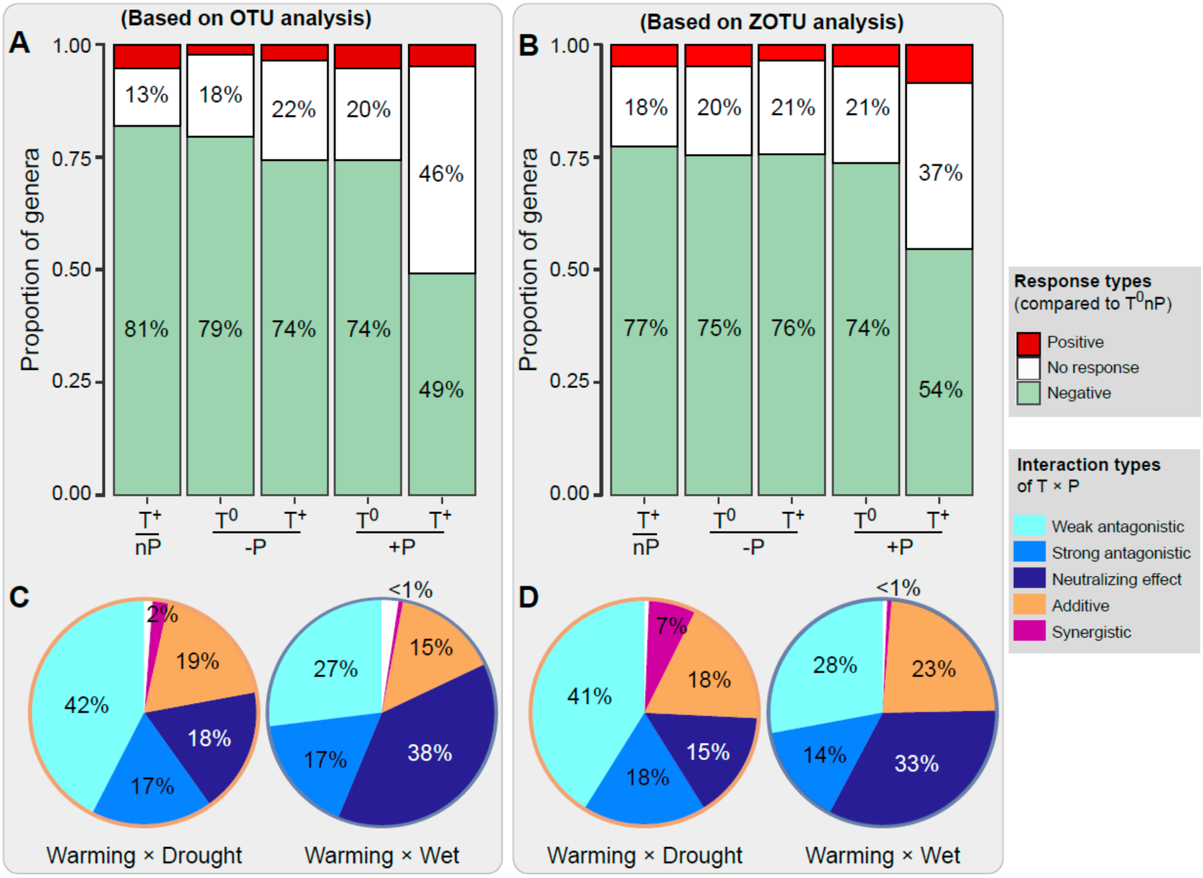


**Fig. S2.**

**The growth responses of grassland bacteria at the genus level to warming and altered precipitation based on OTU analysis (A and C) and ZOTU analysis (B and D). A and B** the single and combined factor effects of climate factors on growth in genera, by comparing with those in $T^0$nP. **C and D** the proportions of genera whose growth influenced by different interaction types of T × P. $T^0$-P represents the ambient temperature and decreased precipitation; $T^+$-P represents warming and decreased precipitation; $T^0$cP represents ambient temperature and precipitation; $T^+$cP represents warming and ambient precipitation; $T^0$+P represents ambient temperature and enhanced precipitation; $T^+$+P represents warming and enhanced precipitation.

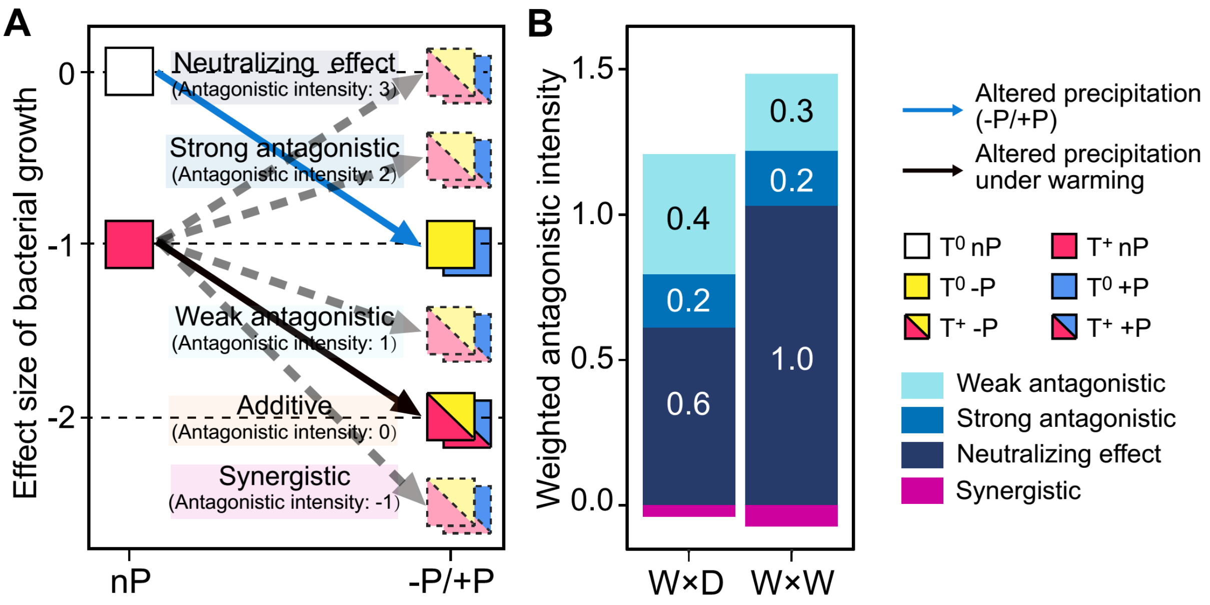


**Fig. S3.**

**The higher level of antagonism of wet × warming than that of drought × warming. A** The antagonistic intensities were assigned to the five interaction types on a 5-point scale, from -1 to 3 for synergistic, additive, weak antagonistic, strong antagonistic and nullifying effect, respectively. **B** The overall antagonistic intensities of all incorporators under warming × drought and warming × wet scenarios were estimated by weighting the relative proportions of incorporators subjected to different interaction types. $T^0$-P represents the ambient temperature and decreased precipitation; $T^+$-P represents warming and decreased precipitation; $T^0$cP represents ambient temperature and precipitation; $T^+$cP represents warming and ambient precipitation; $T^0$+P represents ambient temperature and enhanced precipitation; $T^+$+P represents warming and enhanced precipitation.

**Table S1 The nearest taxon index (NTI) for incorporators subjected to different interaction types under two climate change scenarios.**

| Climate change scenarios | Interaction types | NTI value | *P* value |
|---|---|---|---|
| Warming × Drought | Weak antagonistic | 1.75 | **0.036** |
| | Strong antagonistic | 3.06 | **0.003** |
| | Neutralizing effect | 2.56 | **0.005** |
| | Additive | 2.33 | **0.01** |
| | Synergistic | -0.88 | 0.809 |
| Warming × Wet | Weak antagonistic | 1.49 | 0.063 |
| | Strong antagonistic | 2.08 | **0.021** |
| | Neutralizing effect | 1.85 | **0.036** |
| | Additive | 1.16 | 0.116 |
| | Synergistic | -0.67 | 0.761 |

*P*-value based on comparison of phylogenetic distance observed and that based on 1,000 permutations of a null model. Values in bold: $P < 0.05$. Low *P*-values (and positive indices) indicate that species are more phylogenetically related than expected by chance (phylogenetically clustering). $T^0$-P represents the ambient temperature and decreased precipitation; $T^+$-P represents warming and decreased precipitation; $T^0$cP represents ambient temperature and precipitation; $T^+$cP represents warming and ambient precipitation; $T^0$+P represents ambient temperature and enhanced precipitation; $T^+$+P represents warming and enhanced precipitation.
